# A mathematical framework for analyzing wild tomato root architecture

**DOI:** 10.1101/2021.08.12.456185

**Authors:** Arjun Chandrasekhar, Magdalena M Julkowska

**Affiliations:** Department of Computer Science, University of Pittsburgh; Boyce-Thompson Institute; Center for Desert Agriculture, King Abdullah University for Science and Technology, Jeddah, Saudi Arabia

## Abstract

The root architecture of wild tomato, *Solanum pimpinellifolium*, can be viewed as a network connecting the main root to various lateral roots. Several constraints have been proposed on the structure of such biological networks, including minimizing the total amount of wire necessary for constructing the root architecture (wiring cost), and minimizing the distances (and by extension, resource transport time) between the base of the main root and the lateral roots (conduction delay). For a given set of lateral root tip locations, these two objectives compete with each other — optimizing one results in poorer performance on the other — raising the question how well *S. pimpinellifolium* root architectures balance this network design trade-off in a distributed manner. Here, we describe how well *S. pimpinellifolium* roots resolve this trade-off using the theory of Pareto optimality. We describe a mathematical model for characterizing the network structure and design trade-offs governing the structure of *S. pimpinellifolium* root architecture. We demonstrate that *S. pimpinellifolium* arbors construct architectures that are more optimal than would be expected by chance. Finally, we use this framework to quantify structural differences between arbors grown in the presence of salt stress, classify arbors into four distinct architectural ideotypes, and test for heritability of variation in root architecture structure.

## Introduction

Root architecture in plants is driven by the development of lateral roots that branch off of the main root [1]. Our goal is to derive a model that explains the principles governing branching patterns in *S. pimpinellifolium* root architecture. Root architecture design that affords efficient resource transport is crucial to the overall productivity of the plant [2]; however, it can be advantageous to minimize the amount of material required to construct arbors due to environmental constraints such as resource limitations [3], soil compaction [4], growth environment orientation [5], and salt stress [6]. Prior work models plant arbors as a weighted graph, and uses the theory of Pareto optimality to resolve the trade-off between minimizing resource transport delay and minimizing material cost [7, 8, 9]. We extend prior work by applying a model developed for *Arabidopsis* to quantify design trade-offs in *S. pimpinellifolium*.

Root architecture is highly plastic, showing adaptability to the environments and stress conditions such as salt stress [10]. These salt-induced changes in root architecture were previously used to identify genetic components contributing to overall stress-resilience [6]. Through screening *S. pimpinellifolium*, which is stress-resilient relative to cultivated tomato, we aim to identify novel strategies that contribute to *S. pimpinellifolium* resilience. Through modeling and analyzing network topology, we attempt to gain further insight into network efficiency and adaptability to stress conditions.

In addition to observing high stress-resilience, we find that *S. pimpinellifolium* arbors visually classified into four distinct architectural ideotypes: “Christmas Tree”, “Telephone Pole”, “Droopy Telephone Pole”, and “Broomstick” (Fig 1A). However, a preliminary analysis revealed that this classification scheme is not associated with root structure variation according to traditional physical features (Fig 1B–D). Our goal was to study alternate methods for quantifying network topology, to test whether our qualitative classification scheme captures meaningful variation in root architecture.

**Figure 1:**
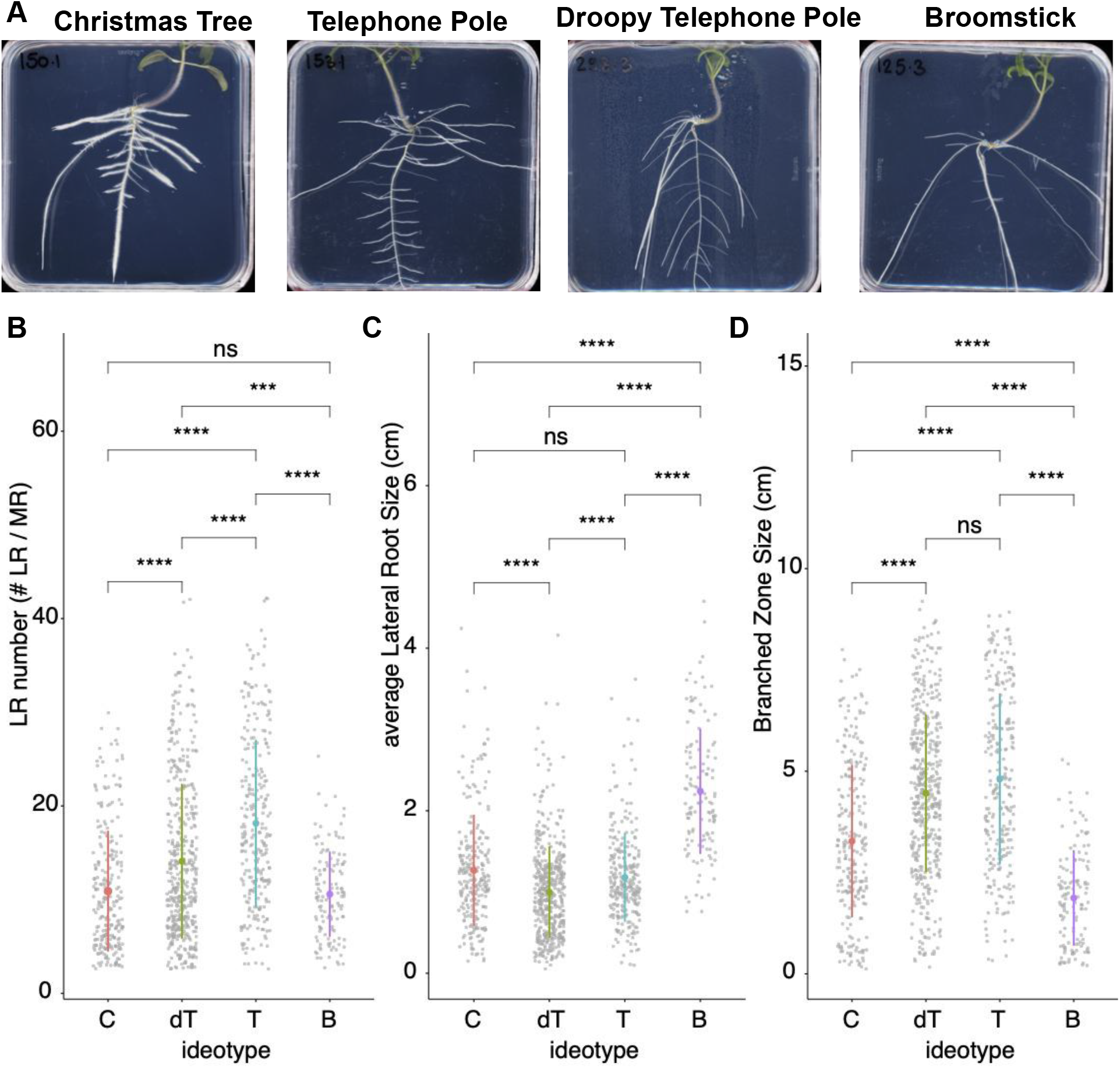
*S. pimpinellifolium* root ideotypes across the diversity panel. A) The four ideotypes of root system architecture were identified in representative accessions: M150 / LA1686 for Christmas Tree; M153 / LA1689 for telephone pole; M125 / LA1628 for broomstick; M298 / LA3160 for droopy telephone pole. The pictures represent the 12 days old seedling root system architecture grown under non-stress conditions. We compared *S. pimpinellifolium* ideotypes for B) number of lateral roots, C) average lateral root length and D) length of the branched zone. The individual ideotypes are Christmas tree (C), Telephone Pole (T), Droopy Telephone Pole (dT) and Broomstick (B). The asterisks indicate significant differences between pairs of ideotypes using Tukey’s honest significance test. In all cases, at least one pair of ideotypes is not detected as significantly different from each other.

Our manuscript is organized as follows. First, we define a graph-theoretic model *S. pimpinellifolium* arbors that formalizes the objectives of wiring cost and conduction delay. We demonstrate how to resolve the trade-off between these two objectives using the theory of Pareto optimality. We use this framework to investigate how well *S. pimpinellifolium* arbors optimize wiring cost and conduction delay, and infer whether these objectives are significant drivers of root architecture topology. We then investigate variation in how arbors prioritize objectives. We test whether arbors prioritize objectives in the presence of a salt stressor; we propose a scheme for classifying arbors into four architectural ideotypes, and use network design trade-offs to validate this classification scheme; finally, we explore whether genetic variation is associated with arbors prioritizing objectives differently.

### Related work

In modeling and analyzing the network topology of *S. pimpinellifolium*, we relate our work to the Euclidean Steiner tree optimization problem [11, 12, 13]. We study the Steiner tree problem with certain constraints on the solution space; additionally our Steiner trees must balance two cost functions instead of one.

Our work is also closely related to the problem of finding a Light Approximate Shortest-Path (LAST) Tree [14, 15]. Given a rooted graph, the goal is to find a spanning tree whose total weight is not too high while maintaining short paths from the root to each vertex. Our work attempts to solve a similar problem but allows for branch points that are not part of the original network.

## Materials and Methods

Below we describe how we use graph theory, as well as the principle of Pareto optimality, to model *S. pimpinellifolium* root architecture and quantify design trade-offs.

### A graph-theoretic framework for studying *S. pimpinellifolium* architecture

We model the *S. pimpinellifolium* root architecture as a weighted, connected, acyclic graph *A* = (*V, E*). Each of the vertices *v* ∈ *V* represents the 3-dimensional coordinate of one of the points on the root. We partition the vertices into two disjoint sets: The *main root vertices R* ⊂ *V*, and the *lateral root vertices L = V* \*R*.

The main root vertices from a path graph – that is, they can be ordered *R* = {*r*_1_, *r*_2_,…, *r*_*n*_} such that (*r*_*i*_, *r*_*i*+1_) ∈ *E* for all 1 ≤ *i* < *n*. We designate *r*_1_ as the *main root base*.

Each lateral root vertex has degree 1, and is directly connected to one of the main root vertices. Lateral root vertices do not connect to another lateral root segment.

For each edge (*u, v*) ∈ *E*, let the *weight w*(*u, v*) be the Euclidean distance between *u* and *v*. For each pair of points *u, ∈ v V*, let the *distance d*(*u, v*) be the length of the (unique) path from *u* to *v*.

Let the *wiring cost* be the sum of the edge lengths of all edges in the graph:

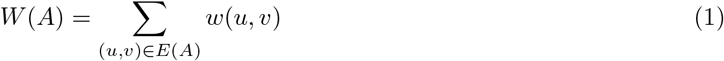

Let the *conduction delay* be the sum of graph distances from the main root base to each lateral root:

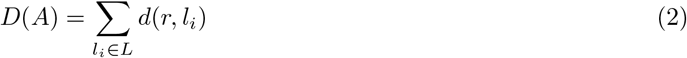

### Pareto-optimal root architecture problem

In general, it is impossible to design a root architecture that simultaneously minimizes wiring cost and conduction delay. To resolve this trade-off, we turn to the principle of Pareto optimality [16] to define the problem of constructing an optimal arbor.

As input, we are given the following (Figure 2A):

- A set of main root points {*r*_1_, *r*_2_,…, *r*_*n*_} that are all connected to each other; in other words, our network starts with the edges {(*r*_1_, *r*_2_), (*r*_2_, *r*_3_),…, (*r*_*n*–1_, *r*_*n*_)}
- A set of lateral root tips {*l*_1_, *l*_2_,…, *l*_*m*_}. Initially these points are not connected to the main root.
- A value *α* ∈ [0, 1]

**Figure 2:**
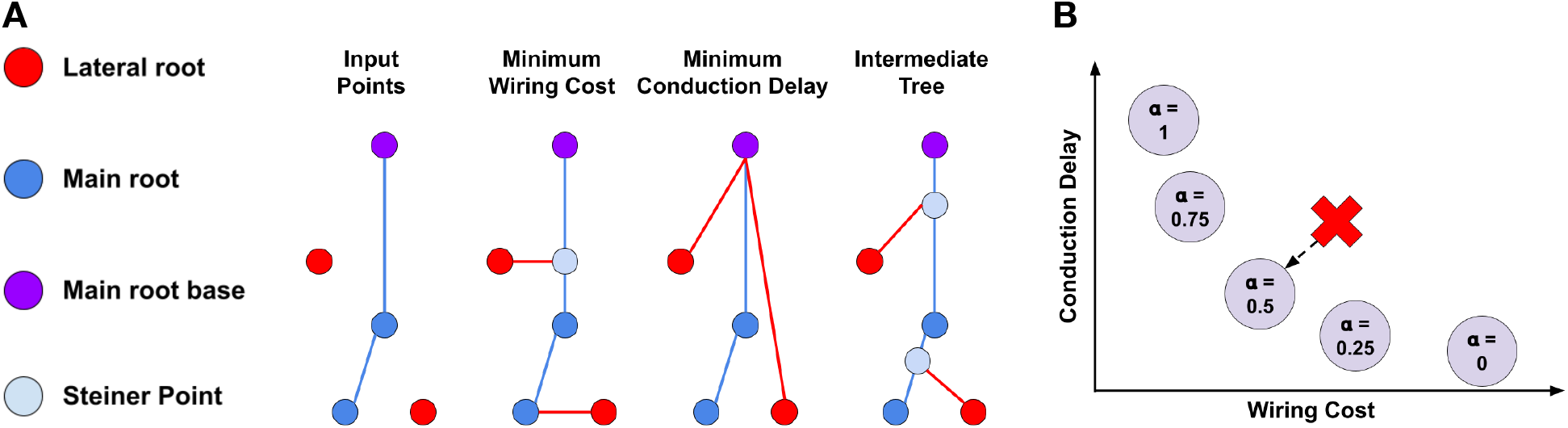
Pareto front of optimal solutions. A) Initially we are given a set of line segments comprising the main root, as well as several lateral root tips that must be connected to the main root. Three possible optimal architectures are shown: one which minimizes the total material cost, one which minimizes the time to transport resources to the main root base, and one which attempts to strike a balance between the two objectives. B) The *Pareto front* of optimal solutions. For a given set of points, we vary *α* ∈ [0, 1] to compute the set of architectures that achieve an optimal trade-off between the two objectives. The red ‘X’ denotes the costs of an observed arbor; we can measure how close the arbor was to being Pareto-optimal, and the Pareto-optimal tree to which the observed arbor was most similar.

We then construct an architecture graph *A* by connecting each lateral root to the main root (Figure 2A). Each lateral root may connect to one of the initial main root points, or it may connect to a midpoint on one of the edges (*r*_*i*_, *r*_*i*+1_) (thus splitting that edge into two smaller edges). The architecture graph *A* should minimize:

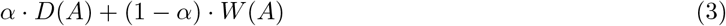

The parameter *α* determines how much the solution should prioritize one objective over the other. Different choices of *α* will yield different optimal solutions *A*_*α*_ that the arbor could have constructed (Figure 2A). We call these optimal architectures the *Pareto front of optimal solutions*.

### An algorithm for generating optimal root architectures

To generate optimal root architectures, we first note lateral roots must connect directly to the main root (and not to each other); thus we can optimize the full architecture by optimizing each lateral root’s connection point independently. To do so, we discretize the main root by dividing each line segment into 100 smaller line segments, thus creating several new potential connection points. For each lateral root, we consider which of connection point on the main root is optimal for that lateral root according to equation (3).

Note that this algorithm is an approximation, and can be made exact using calculus; such improvements are the subject of ongoing future work.

### Quantifying network design trade-offs in observed architectures

Given an observed architecture *A*, we may use the above algorithm to compute the set of optimal root architectures: {*A*_*α*_│*α* ∈ [0, 1]} (Figure 2A–B). Using this, we can calculate two metrics for quantifying network design trade-offs in the original observed arbor (Figure 2B):

- **Pareto front distance**: The L-2 distance between costs of arbor *A* and the costs of the closest optimal tree:

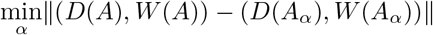 This measures how close the original arbor came to managing an optimal trade-off between wiring cost and conduction delay.
- **Pareto front location**: The value of *α* ∈ [0, 1] associated with the optimal arbor closest in cost to the observed arbor.

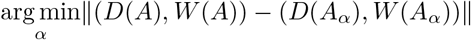 The closer this value to 1, the more strongly the arbor prioritized minimizing wiring cost at the expense of conduction delay.

Under this framework: we tested four hypotheses: 1) *S. pimpinellifolium* roots are closer to being Pareto optimal than would be expected by chance; 2) *S. pimpinellifolium* roots respond to salt stress by modifying network design trade-offs; 3) variation in Ideotype classification is associated with variation in Pareto front location; 4) Pareto front location is a heritable trait that may be driven (in part) by genetic factors.

### Comparing observed arbors against a null model

To evaluate how well *S. pimpinellifolium* would be expected to optimize wiring cost and conduction delay by chance, we compared observed arbors with arbors that were randomly generated on the same set of points. To generate a random arbor, we start with the same main root segments, and same lateral root tip locations, as the original observed arbor. Each lateral root tip is then connected to a point chosen at uniform random from the set of line segments comprising the main root.

## Empirical Data

The natural diversity panel 260 accessions of *S. pimpinellifolium* was screened for salt-stress induced changes in root system architecture. The seeds were surface sterilized using 50% bleach for 10 minutes and washed 4-6 times using sterile MiliQ water. The seeds were stratified at 4°C for 24 hours to improve germination. The seeds germinated on square petridishes (12 x 12 cm) on 1/4 Murashi-Skoog media (1.1 g MS, 5 g sucrose, 1 g M.E.S. buffer, pH 5.8 KOH, 10 g Dashin agar). Four days after germination, the seedlings were transferred to square petridishes containing 1/4 Murashi-Skoog media for Control treatment or 1/4 MS media supplemented with 100 mM NaCl for Salt treatment. The plates were scanned every day for 7 consecutive days.

The root system architecture was quantified from the images recorded 0, 1, 2, 3 and 4 days after transfer to Control and Salt plates using SmartRoot [17]. The global root data was compiled and analyzed in R for lengths of main and lateral roots as well as growth rates of main root, and lateral roots. The root node data was used for calculating the Pareto optimality.

Plants were manually and qualitatively classified into one of four representative ideotype accessions (Figure 1A). The root systems where lateral roots were decreasing in length throughout the main root axis were classified as “Christmas Tree”. The root systems where the lateral root was constant throughout the main root axis and where lateral roots were growing at approximately 90°angle from the main root axis were classified as “Telephone Pole”. The root systems similar to “Telephone Pole”, but with lateral roots growing under approximately 45°angle from the main root axis were classified as “Droopy Telephone Pole”. The root systems where majority of the lateral roots originated at the root base were classified into “Broomstick” ideotype.

## Results

### *S. pimpinellifolium* roots are Pareto optimal

For each arbor, we computed the Pareto front of optimal architectures that the arbor could have constructed, and measured each arbor’s distance to its respective Pareto front. To test whether arbors are closer to being Pareto optimal than would be expected by chance, we compared each arbor against 20 randomly generated arbors (Figure 3A). We find that *S. pimpinellifolium* arbors are more optimal than random trees 98.61% of the time, which is significantly higher than 50% (binomial test, *p* < 10^−322^). On average, the *S. pimpinellifolium* arbor is 3.15 ± 2.60 times closer to the Pareto front than the random arbor; this ratio is significantly higher than 1 (Welch’s T-test; *p* < 10^−322^). We conclude that *S. pimpinellifolium* architecture optimizes wiring cost and conduction delay significantly more than would be expected by chance. This suggests that further study of *S. pimpinellifolium* may yield new insights into design of distributed network optimization algorithms.

**Figure 3:**
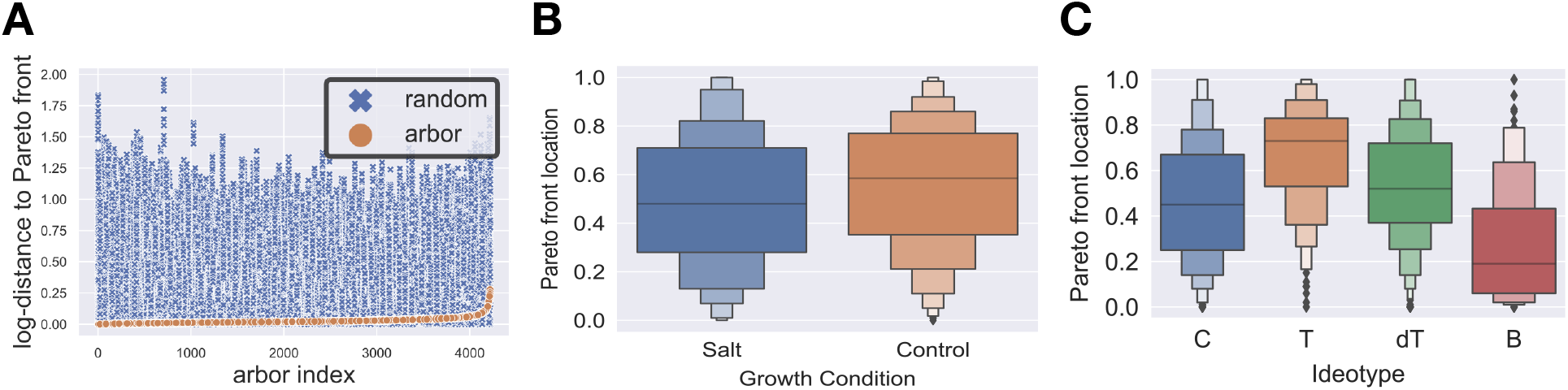
Analysis of Pareto front distance and location. A) For each *S. pimpinellifolium* arbor, we calculated the distance to the Pareto front, as well as the distance for 20 arbors that were randomly generated on the same set of main root and lateral root points. We find that *S. pimpinellifolium* are significantly closer to being Pareto optimal than would be expected by chance. B–C) For each arbor, we calculated its closest location on the Pareto front, a measure of how the arbor prioritized objectives. We find significant differences between how arbors grown in different conditions (B), and arbors from different ideotype classes (C), prioritize objectives.

### Salt stressor affects network design trade-offs

Arbors grown in control conditions tended to prioritize optimizing wiring cost more than arbors grown in conditions with a salt stressor (Figure 3B). The average value of *α* for arbors grown in control conditions was was 0.55 ± 0.26, as compared to 0.50 ± 0.28; the difference between the two groups was statistically significant (Mann-Whitney test; *p* < 0.001). This suggests that salt stressors may lead arbors to place extra emphasis on fast resource transport.

### Classification of ideotypes is associated with variation in network design trade-offs

Finally, we find a strong association between an arbor’s architectural ideotype and the degree to which an arbor prioritizes trade-offs between wiring cost and conduction delay (Figure 3C). There are significant differences in Pareto front location between different ideotype groups (Kruskal-Wallis one way ANOVA; *p* < 10^−28^). The Pareto front location was significantly different between all pairs of ideotype groups (Mann-Whitney test with a Bonferroni correction, *p* < 0.01 in all cases). Thus, Pareto front location represents a quantitative measure that is associated with the architectural diversity that we observe upon visual inspection of *S. pimpinellifolium* arbors.

### Network design trade-offs are heritable

The variation for Pareto front location was lower within a single genotype than the variation observed across individual genotypes. We calculated broad-sense heritability 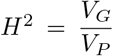, defined as proportion of variation due to genotype specific effects *V_G_* to the observed phenotypic variation across the genotypes *V_P_*; values close to 1 indicate high heritability, and values close to 0 indicate low heritability [18]. Broad-sense heritability for Pareto front location was 0.6345 for root systems measured 4 days after transfer, indicating the high heritability of this trait and its potential use in forward genetic studies.

## Discussion

Like many natural and engineered systems, *S. pimpinellifolium* root architecture must balance trade-offs between multiple competing objectives. Generally there is not a single architecture that is optimal for all possible objectives. Instead, the best the arbor can do is converge to an architecture that is Pareto optimal – meaning it is impossible to improve one objective without sacrificing performance in the other objectives. Our work proposes two objectives – minimizing wiring cost, and minimizing conduction delay – that *S. pimpinellifolium* roots seek to optimize in a Pareto optimal manner. Using graph theory and the principle of Pareto optimality, we define a framework for quantifying the optimality of a given *S. pimpinellifolium* root architecture. We apply this model to *S. pimpinellifolium* imaging data and find that *S. pimpinellifolium* arbors are significantly closer to being Pareto optimal for these objectives than would be expected by chance. We use Pareto front location to show that *S. pimpinellifolium* arbors prioritize objectives differently in the presence of a salt stressor; we find that unlike traditional network design metrics, Pareto front location quantifies *S. pimpinellifolium* architecture in a way that is strongly associated with the architectural diversity we observe qualitatively; and we find that Pareto front location shows a strong association with genetic variation. Overall, our work provides a framework for quantifying *S. pimpinellifolium* based on how and how well they manage network design trade-offs. More broadly, our work provides a blueprint for quantifying how natural and engineered systems manage trade-offs between several competing objectives.

### Future directions

Pareto front location is associated with our classification of *S. pimpinellifolium* arbors into architectural ideotypes. Future work involves using clustering analysis of Pareto front location to conclusively validate our chosen classification scheme (or identify the correct number of clusters). If we find evidence for the existence of our four identified ideotypes, future work will seek to find the functional basis behind each of these four structures.

In addition to formulating an exact algorithm for generating Pareto-optimal arbors, we seek to study the development process of *S. pimpinellifolium*. We will take advantage of data tracking the growth of individual arbors over time to measure how arbors prioritization of objectives changes over time, and eventually define a model for the distributed growth process used to generate Pareto optimal architectures.

Future work includes Genome Wide Association Study (GWAS), to identify underlying genetic components of Pareto front location and distance in *S. pimpinellifolium* roots. Through reverse genetic approaches we aim to further study genetic components of Pareto front location, their role in root architecture development and further developing better understanding on how *S. pimpinellifolium* plants prioritize objectives during non-stress and stress conditions.

Finally, we will expand our model to study the trade-off between not only wiring and transport efficiency, but also gravitropism and growth media exploration. Certain arbors may have a higher tendency to grow towards or away from gravitational forces in the growth medium. Furthermore, certain arbors may exhibit higher nutrient uptake at the main root, thus leading lateral roots to explore farther horizontally in order to avoid competing with the main root. We will expand our mathematical model to include these factors as extra constraints on how lateral roots may connect to the main root (while still optimizing wiring cost and conduction delay.

## Acknowledgements

We thank the organizing committee for the 2021 Biological Distributed Algorithms (BDA) conference, as well as the reviewers for or BDA submission. We thank Guillame Bauchet for help with understanding the output from the plant imaging software. We thank Graham Zug, who is currently working with Dr. Chandrasekhar on mathematical optimization algorithms for constructing Pareto optimal plant arbors.

## Author Confirmation Statement

AC: data curation, formal analysis, methodology, software, validation, visualization, writing (original draft preparation). MJ: conceptualization, data curation, formal analysis, investigation, methodology, visualization, writing (original draft preparation).

## Data availability

We will make data available upon request. Our code for analyzing arbors and performing statistical analysis can be found here https://github.com/arjunc12/Plant-Architecture.

